# Proteome-Driven Phenotyping of Identified Single Neurons in Intact Brain Tissue by Aspiration Patch Proteomics

**DOI:** 10.64898/2026.04.22.720006

**Authors:** Cole C. Johnson, Sam B. Choi, Juan Zegers-Delgado, Alexandre Kisner, Ricardo C. Araneda, Abigail M. Polter, Peter Nemes

## Abstract

Single-cell proteomics has advanced rapidly, but direct proteome measurements from identified neurons in intact brain tissue remain difficult because most workflows require cell isolation and recent patch-based studies have emphasized whole-soma retrieval. Here we show that aspiration-based patch proteomics enables deep proteome profiling of identified single neurons directly in acute mouse brain slices. We combined fluorescence-guided patch-clamp microsampling, minimal-loss bottom-up proteomics, and high-sensitivity capillary electrophoresis–timsTOF mass spectrometry to analyze partial somal aspirates from dopaminergic, parvalbumin, and serotonergic neurons *in situ*. The workflow identified more than 1,000 proteins from single-neuron samples under optimized conditions while consuming only about 0.25% of the processed digest per analysis. These proteomes were sufficient to separate biological replicates by neuronal phenotype, distinguish neuronal subtypes on the basis of protein expression alone, and define a conserved somal proteome shared across neuronal classes. Our results establish that controlled aspiration of partial somal material can support proteome-driven phenotyping without whole-soma retrieval, cell dissociation, or loss of native tissue context. Aspiration patch proteomics therefore provides an accessible route for subtype-level proteome phenotyping in intact brain tissue.

## INTRODUCTION

Neuronal phenotypes are shaped by proteomic programs that influence excitability, synaptic transmission, metabolism, and plasticity.^1-3^ Single-cell transcriptomics has exposed extensive neuronal diversity,^4-7^ but transcript abundance is an imperfect proxy for protein composition because translation, turnover, and activity-dependent regulation decouple RNA and protein levels. Direct single-neuron proteomics in native tissue therefore remains an important and still underdeveloped analytical target.

Recent advances in mass spectrometry (MS) have pushed discovery proteomics to the single-cell scale (reviewed in ^8-10^). Improvements in sample preparation, separations, and MS sequencing have enabled up to ∼6,500 proteins to be detected from isolated single cells.^11,12^ That depth is impressive, but the standard route still depends on prior cell isolation and therefore strips away spatial context and any direct link to physiology. Capillary microsampling has partly solved that problem for tissue-bound cells. When paired with capillary electrophoresis (CE), it enabled detection of up to ∼1,700 proteins from individual blastomeres in developing *Xenopus laevis* and zebrafish embryos and revealed cellular heterogeneity that would have been masked in pooled measurements.^13,14^ Recent electrophoresis-correlative (Eco) acquisition strategies have further improved CE–MS depth.^15,16^ Neurons in mammalian brain tissue remain particularly difficult targets, however, because they are chemically complex, physically small, and embedded in a crowded cellular environment that challenges low-input proteomics.

Patch-clamp electrophysiology provides direct physical access to somal contents through the recording pipette and therefore offers an obvious entry point for single-cell molecular analysis. Patch-Seq proved how valuable that access can be for transcriptomics in intact tissue.^17-20^ We previously adapted the same logic to aspiration-based patch proteomics coupled to CE–MS and detected 175 proteins from about 1 pg of aspirated material, corresponding to ∼0.25% of the somal proteome of a single dopaminergic neuron.^21,22^ Since then, patch-based studies that physically retrieved whole somata or whole neurons have reported deeper coverage in culture and acute slices.^14,23-25^ Those studies expanded the feasible depth range, but they are not directly equivalent to aspiration-based patch proteomics because they recover substantially more material and, in the most recent acute-slice work, were implemented on newer-generation Orbitrap instrumentation. Reported depth therefore depends on sampling format, recovered material, instrument platform and the fraction of digest that is ultimately analyzed.

We therefore asked whether aspiration-based, MS-compatible patch microsampling could reproducibly deliver proteomes of sufficient depth and quality for subtype-level phenotyping directly in intact mammalian brain tissue. Specifically, we tested whether internal-solution chemistry could be tuned to balance electrophysiology with proteomics, whether aspirated somal material could support deep CE–timsTOF Pro measurements from acute brain slices, and whether these partial somal proteomes were sufficient for subtype-level classification while defining a shared somal core proteome. The study therefore establishes an analytical framework for determining how much phenotypic information can be extracted from controlled partial somal sampling rather than whole-soma retrieval.

## RESULTS 2,500-3,500 words

Our workflow integrated fluorescence-guided patch microsampling with minimal-loss bottom-up proteomics and CE–timsTOF Pro MS for single-neuron analysis in acute mouse brain slices (**Fig. 1**). The central analytical question was whether controlled aspiration of partial somal material, rather than whole-soma retrieval, could reproducibly provide proteomes of sufficient depth for subtype-level phenotyping in native tissue. We first optimized internal-solution chemistry because this variable simultaneously controls recording quality, compatibility with CE stacking and electrospray performance, and peptide sequencing sensitivity. We then applied the optimized workflow to fluorescently labeled dopaminergic (DA), parvalbumin (PVN), and serotonergic (5-HT) neurons and tested whether their somal proteomes supported unsupervised classification and subtype-resolved quantitative comparisons.

**Figure 1.**
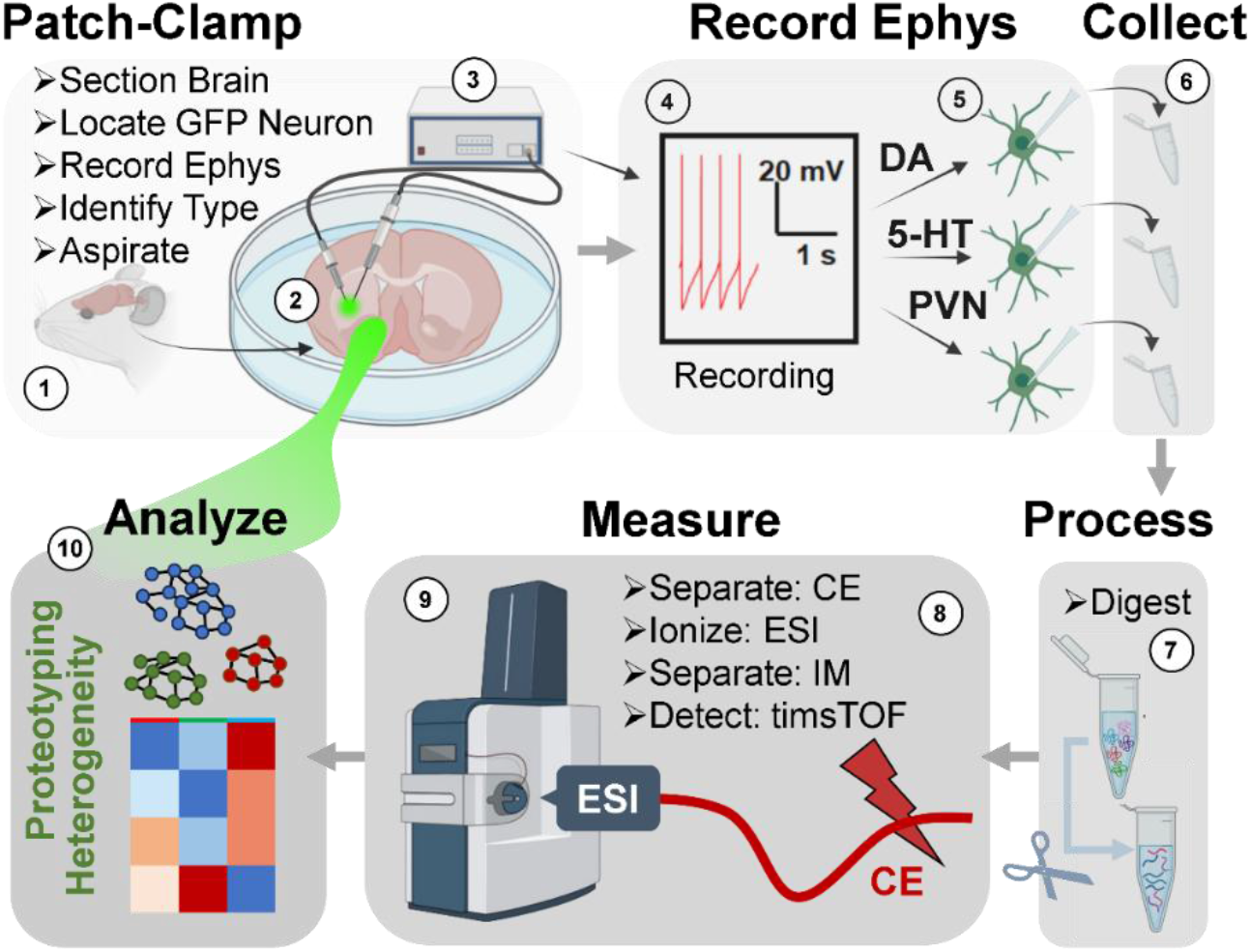
Patch-guided single-neuron proteotyping in acute brain slices by capillary electrophoresis (CE) timsTOF mass spectrometry (MS). A patch pipette was guided under electrophysiological control to the soma of a fluorescently identified neuron in an acute mouse brain slice. Following seal formation and brief recording, somal material was aspirated through the pipette and expelled into low-binding microtubes for minimal-loss bottom-up proteomics rather than by physically retrieving the intact soma. Peptides were separated by high-efficiency CE, ionized by high-sensitivity nanoESI, and detected by timsTOF MS controlled under electrophoresis-correlative data-dependent acquisition. Fluorescence monitoring acquired before and after aspiration confirmed sampling of the targeted soma while leaving neighboring neurons intact. This workflow enabled ultrasensitive proteomic analysis of limited neuronal material in native tissue.

Method development was performed in visually identifiable mitral cells from acute olfactory bulb slices (**Fig. 2**), which offered large somas, reliable access for patch-clamp recording and well-characterized firing behavior.^26-31^ Cell identity was confirmed based on location in the brain and morphology under fluorescence and transmitted light, and characteristic action potential firing patterns during whole-cell current-clamp recordings. Conventional 2 M potassium gluconate supported stable electrophysiology, but proteomic output largely collapsed under these conditions, yielding only 23 cumulative protein identifications across technical duplicates (Supplementary **Table S1**). This result is consistent with the expected effects of a highly concentrated non-volatile salt load undermining field-amplified stacking in CE and suppressing ionization during MS analysis.

**Figure 2.**
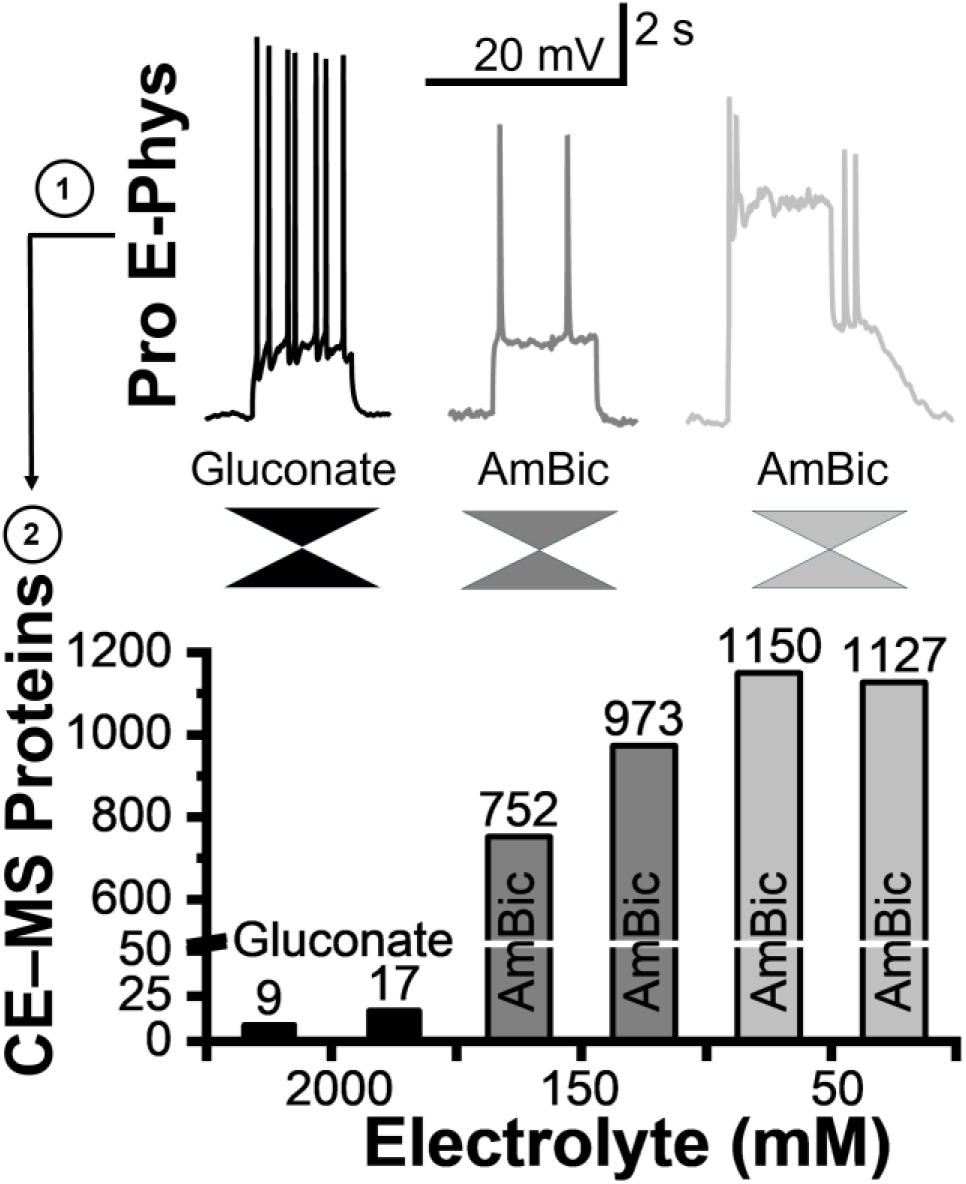
Trade-off between internal electrophysiology and proteome depth. Olfactory bulb neurons were patched using potassium gluconate (2 M) or ammonium bicarbonate (AmBic: 150 mM or 50 mM) internal solutions and ∼0.25% of the processed proteome digest from aspirated somal material was analyzed using CE–timsTOF MS in technical duplicate. Potassium gluconate supported robust firing but suppressed proteomics performance, whereas AmBic-based internal solutions increased proteome coverage but compromised electrical recording and neuron viability.

We therefore evaluated ammonium bicarbonate (AmBic) as a volatile, MS-compatible alternative. The 50 mM AmBic condition produced the deepest cumulative proteome, with 1,353 proteins identified (**Table S1**), but membrane integrity deteriorated too rapidly to support reliable electrophysiology. Increasing the AmBic concentration to 150 mM reduced cumulative identifications to 1,061 proteins but improved osmotic support and recording stability. These results identify internal-solution chemistry as a key analytical determinant of aspiration-based patch proteomics because it governs the trade-off between electrophysiological compatibility and downstream CE–MS performance. On this basis, 150 mM AmBic was selected as the practical operating condition for subsequent experiments.

Using fluorescent reporter lines, we then targeted DA neurons in the ventral tegmental area, PVN interneurons in the prefrontal cortex, and 5-HT neurons in the dorsal raphe nucleus. A standard patch pipette was sealed onto the soma, and somal material was aspirated rather than extracting the entire cell body from the slice. Fluorescence fell sharply in the sampled soma, fluorescence appeared in the pipette and neighboring labeled cells remained visually unchanged, consistent with selective microsampling (**Fig. 3**). Aspirated volumes were estimated at ∼1–10 nL from soma size, fluorescence loss, and pipette-resistance changes. The collected aspirate proteomes were digested with a minimal-loss workflow and analyzed by CE–timsTOF Pro MS, with about 0.25% of the processed digest injected per run.

**Figure 3.**
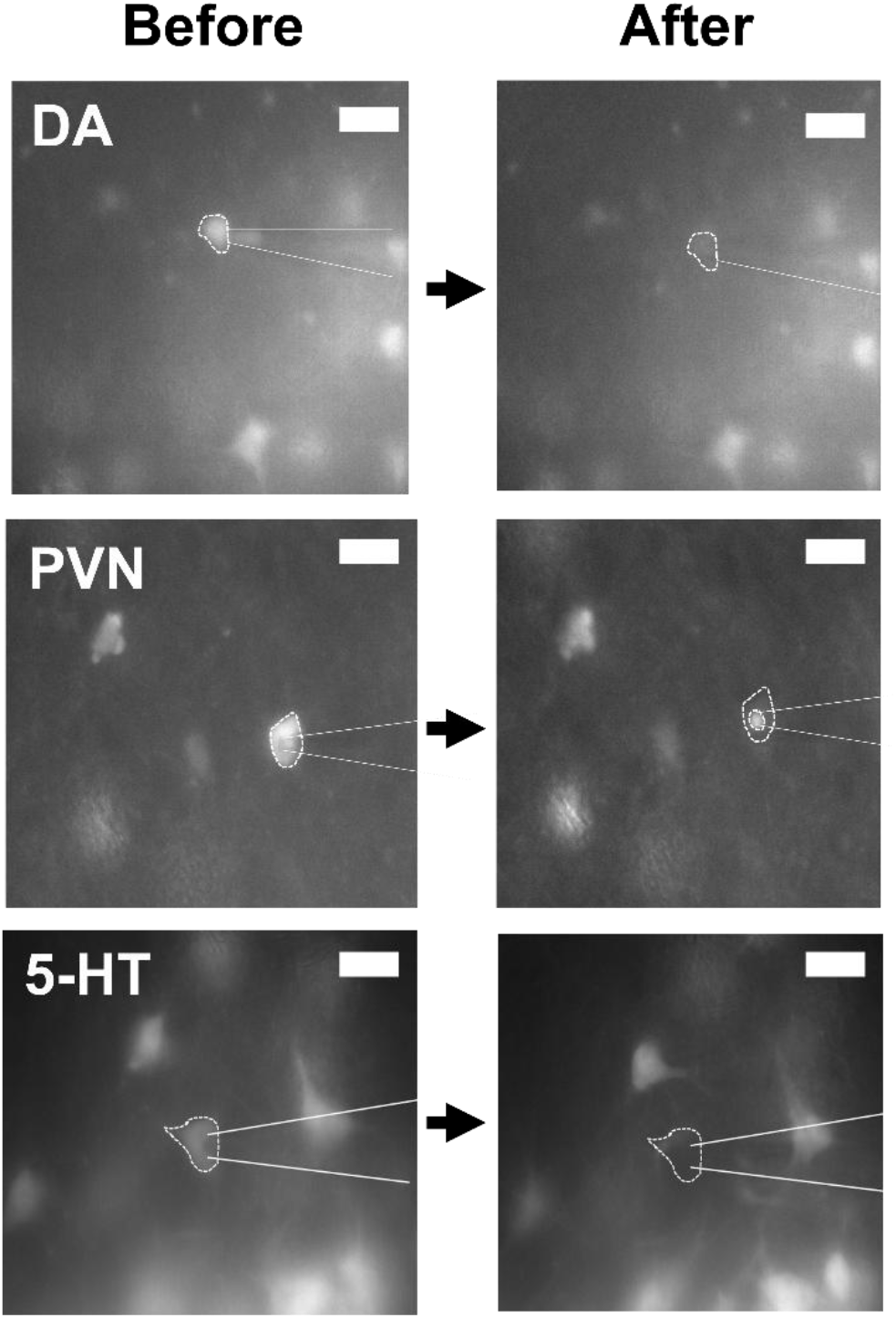
Fluorescence-guided microsampling of individual neuronal somas. Fluorescent dopaminergic (DA), parvalbumin (PVN), and serotonergic (5-HT) neurons were targeted in the ventral tegmental area, prefrontal cortex, and dorsal raphe nucleus under epifluorescence microscopy, respectively. Somal material was aspirated into a patch pipette without physical retrieval of the intact soma. Fluorescence decreased in the sampled soma after aspiration, consistent with microsampling of the somal content.

Across the somal aspirates, 1,894 unique protein entries were identified, with cumulative counts of 1,798 in the DA, 1,248 in the PVN, and 1,428 in the 5-HT neurons (**Fig. 4A**; **Table S2**). The identified proteomes were consistent with expected somal composition and included metabolic, organellar, signaling, and canonical neuronal structural proteins such as microtubule-associated protein 2 (MAP2), tau, and neurofilament heavy, medium, and light polypeptides. Because GFAP is classically associated with astrocytes, its presence may reflect minor contamination from adjacent glial processes during microsampling, although neuronal GFAP expression has also been reported under stress or pathology.^32,33^ Even with this caveat, the data show that limited somal aspirates can support substantially deeper proteome coverage than our earlier aspiration-based implementation, thereby extending the measurable range of partial-sampling patch proteomics.

**Figure 4.**
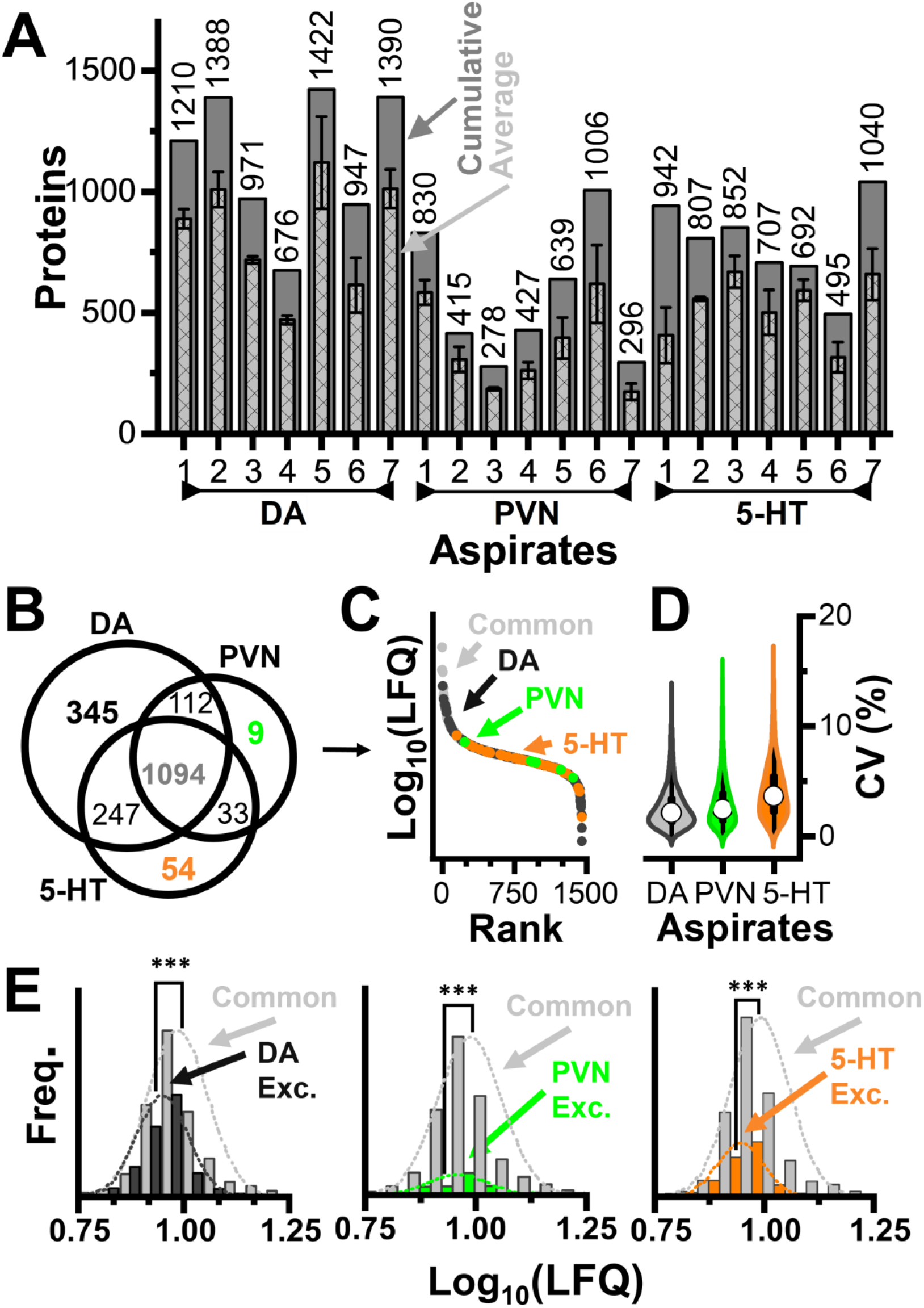
Proteome depth and quantitative performance. Somal aspirates from DA, PVN, and 5-HT neurons were analyzed by CE–timsTOF MS with n = 7 biological replicates per neuronal class and technical triplicates for each sample. **(A)** Proteome depth across biological replicates, shown as per-cell identifications with average and cumulative counts. **(B)** Overlap of identified protein groups across neuronal classes. **(C)** Distributions of label-free quantitative (LFQ) intensities across neuron types revealing broad endogenous concentration range. **(D)** Low coefficients of variation (median < 5%) across biological replicates, demonstrating quantitative reproducibility. **(E)** Comparison of mean LFQ abundance distributions for neuron-type-exclusive and shared proteins, indicating that proteins exclusive to each neuron type were generally lower in abundance than those commonly identified. These data were obtained from aspiration-based samples while only ∼0.25% of the processed digest was analyzed per run, highlighting the depth achievable from limited neuronal material. Key: ***, *p* < 5 × 10^−5^ (Mann-Whitney).

A total of 1,703 of these proteins were also quantified among multiple biological replicates and phenotypes. The majority of proteins were shared across neuronal subtypes, with smaller subsets detected preferentially within each class (**Fig. 4B**). The measured label-free quantitative (LFQ) abundances spanned multiple orders of magnitude, consistent with a broad anticipated endogenous concentration range (**Fig. 4C**). Quantification was robust, with Pearson correlation coefficients exceeding ρ = 0.75 and mean coefficients of variation below 5% across the biological replicates within each neuronal phenotype (**Fig. 4D**). Proteins quantified exclusively in one neuron type showed significantly lower abundance distributions comparable to those shared (**Fig. 4E;** *p* = 2.2 × 10^−16^ for DA, 5.6 × 10^−5^ for PVN, and 2.2 × 10^−16^ for 5-HT). Based on STRING reactive pathway analysis, these proteins were enriched in GAP junction trafficking and regulation, reception activation, membrane trafficking, and metabolism of amino acids and derivatives. These measurements established that aspiration-derived somal proteomes were sufficiently reproducible and information-rich to support comparative phenotyping across neuronal subtypes.

Principal component analysis (PCA) of label-free quantification data clustered biological replicates according to neuronal phenotype in the scores plot (**Fig. 5A**). The corresponding loading plot showed that this separation was driven by both shared and variable protein-expression patterns across the measured somal proteomes, with representative examples shown for Sirt2, Marcksl1, and Arl8a. Consistent with the PCA results, hierarchical cluster analysis (HCA) also grouped biological replicates by known cell type, and the heat map of the top 250 differential proteins cleanly separated the three neuronal phenotypes (**Fig. 5B** and **Fig. S1**). In this analysis, PVN and 5-HT somal proteomes clustered more closely to each other than to DA proteomes. Together, these results show that aspiration-derived single-neuron somal proteomes acquired directly from intact brain tissue contained sufficient information for subtype classification without reliance on morphology or electrophysiological signatures.

**Figure 5.**
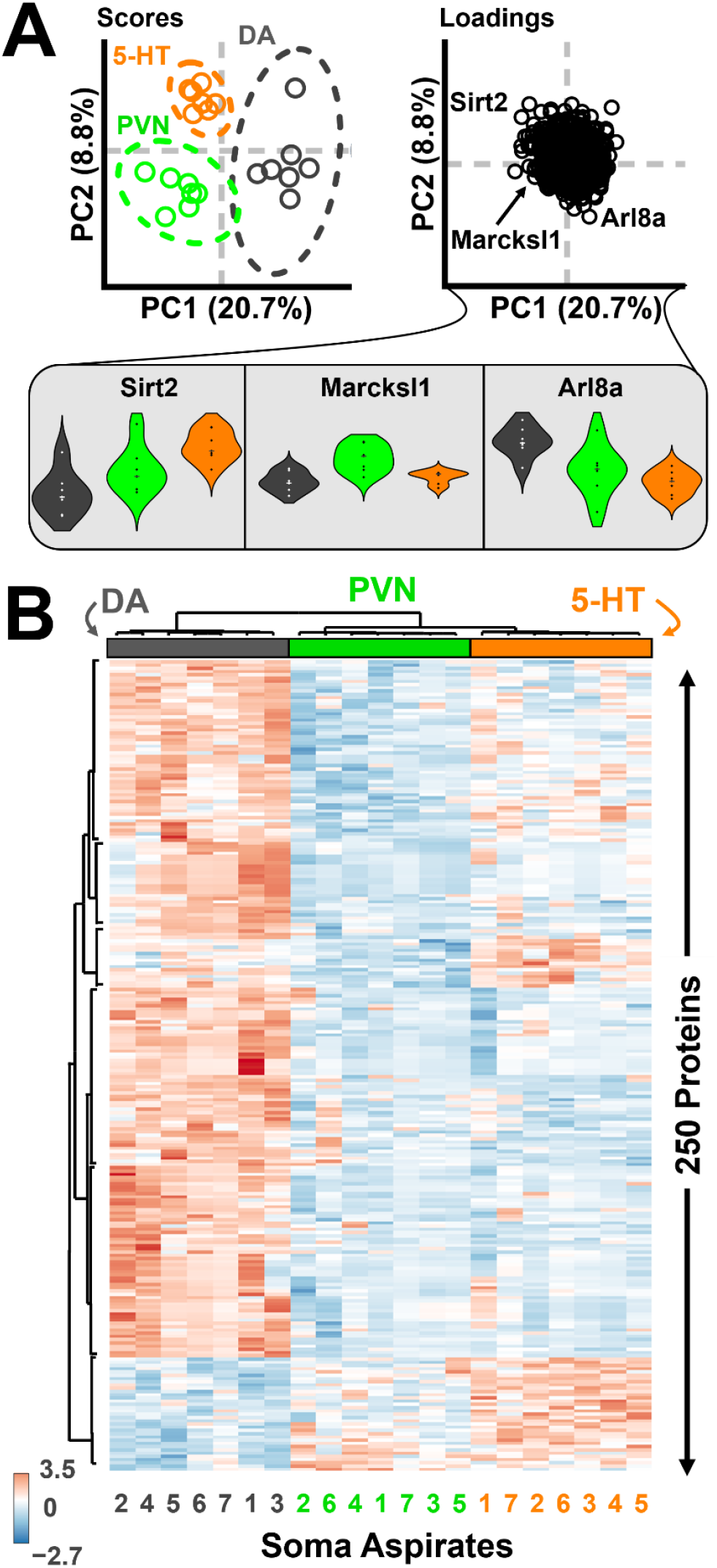
Proteomic signatures classify dopaminergic (DA), parvalbumin (PVN), and serotonergic (5-HT) neurons *in situ*. **(A)** PCA scores plot (left) shows clear clustering of biological replicates for different neuronal subtypes. Ellipses show 95% confidence interval around the biological replicates within each phenotype. PCA loading plot (right) reveals differing patterns of protein expression along principal component axes. Violin plots (bottom insets) show representative protein profiles. **(B)** Hierarchical cluster analysis of the 250 statistically most different features (zoom-in, **Fig. S1**), with heatmap showing z-scored abundances of the top proteins discriminating among subtypes (one-way ANOVA). Individual neurons clustered according to their known subtype identity, indicating that proteomic signatures alone were sufficient to distinguish the neuron types in intact tissue. This analysis established a proof-of-concept for in-tissue single-neuron proteotyping.

To further define protein-level differences between neuronal subtypes, pairwise statistical comparisons were performed on LFQ abundances using the Wilcoxon rank-sum framework and FDR control described in the **Methods**. The DA-versus-PVN contrast produced the largest variable set, the DA-versus-5-HT contrast was intermediate, and the PVN-versus-5-HT contrast was comparatively sparse (**Table S3**). Against that variable minority, **Table S4** defines a conserved somal core proteome of 1,129 proteins with no significant abundance differences (*p* ≥ 0.05) across all pairwise comparisons. STRING analysis organized this stable set into interacting modules associated with actin and cytoskeletal organization, synaptic machinery, mitochondrial energy metabolism, and gene-expression/proteostasis functions (**Fig. 6B**). This dataset defines a stable somal proteomic baseline across neuronal subtypes and supports the interpretation that the differential features resolved elsewhere primarily reflect biological differences among the sampled neuronal subtypes rather than technical instability.

**Figure 6.**
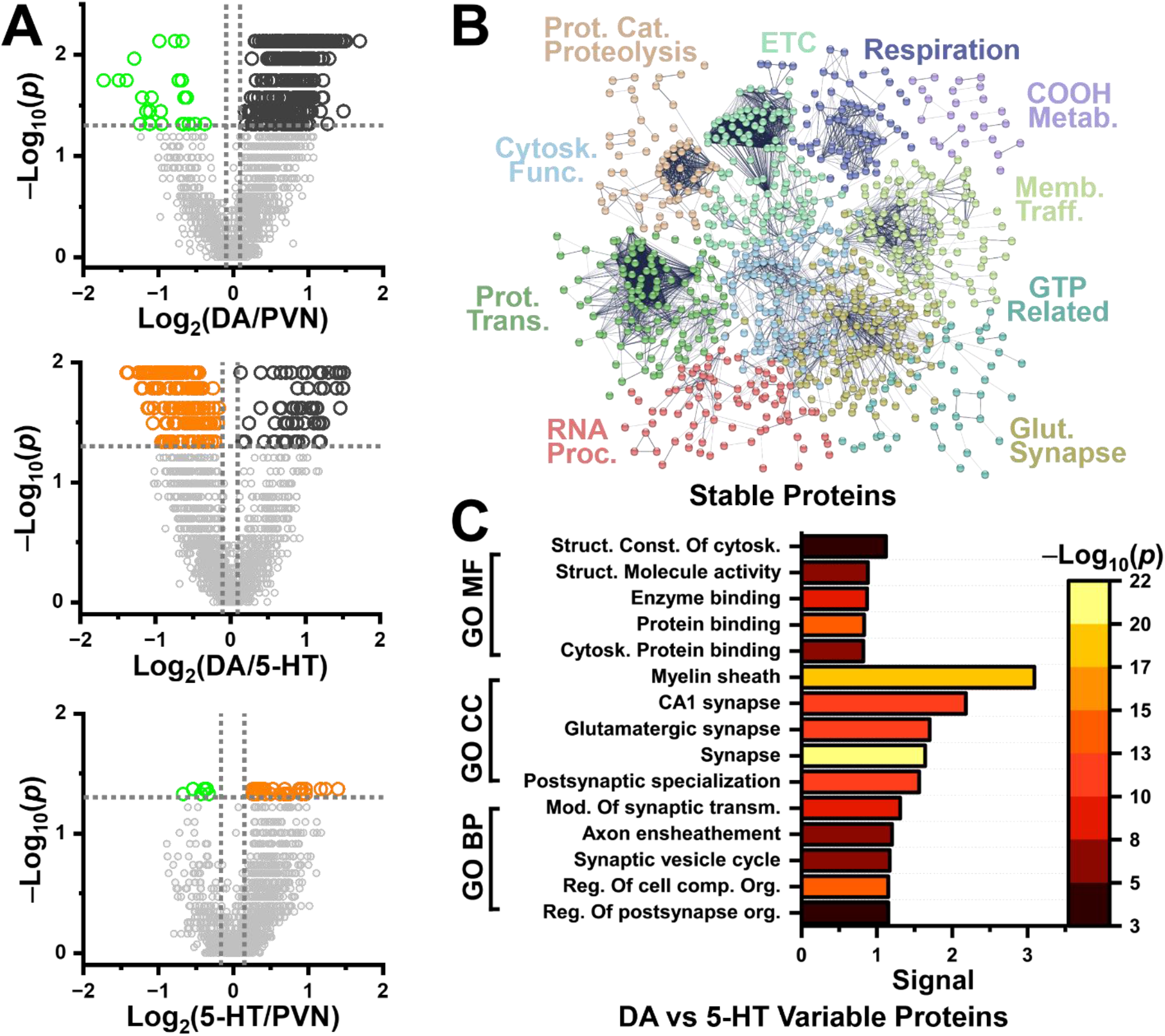
Pairwise statistical analysis defines conserved and subtype-variable somal proteomes. **(A)** Pairwise statistical comparisons of protein abundances between DA, PVN, and 5-HT neuronal somas using the Wilcoxon rank-sum tests for proteins quantified across neuronal classes (**Table S3**), with *p* < 0.05 marking significance (dashed gray lines). **(B)** STRING protein– protein interaction network of proteins with no significant abundance differences across all pairwise comparisons (*p* ≥ 0.05; **Table S4**), defining a conserved somal core proteome. Proteins organized into modules associated with actin dynamics, synaptic machinery, mitochondrial energy metabolism, and gene expression and proteostasis. **(C)** Gene ontology (GO) analysis of significantly different proteins (*p* < 0.05; **Table S5**) across pairwise comparisons. Molecular function (MF) GO terms show significant enrichment of cytoskeletal and structural proteins, while cellular component (CC) and biological process (BP) GO terms are largely neuron- and synapse-specific. Key: Prot. Cat. Proteolysis, proteolysis involved in protein catabolism; ETC, electron transport chain and oxidative phosphorylation; COOH Metab., carboxylic acid metabolic process; Memb. Traff., membrane trafficking; Glut. Synapse, glutamatergic synapse; RNA Proc., RNA processing; Prot. Trans., protein transport; Cytosk. Func., cytoskeletal function; Struct. con. of cytosk., Structural constituent of cytoskeleton; Struct. mol. act., Structural molecule activity; Cytosk. pro. bin., Cytoskeletal protein binding; Schaff. col. CA1 synapse, Schaffer collateral - CA1 synapse; Postsynaptic spec., Postsynaptic specialization; Mod. of chem. syn. transm., Modulation of chemical synaptic transmission; Synapt. ves. cyc., Synaptic vesicle cycle; Reg. of cell. comp. org., Regulation of cellular component organization; Reg. of postsyn. org., Regulation of postsynapse organization.

By contrast, **Table S5** shows that only four variable proteins (*p* < 0.05) were shared across all three pairwise contrasts, whereas most variable proteins were confined to one comparison or shared by only two. This limited overlap is notable given that the measurements were derived from aspirated somal fractions rather than whole-soma retrieval, indicating that even partial sampling of the neuronal proteome is sufficient to resolve subtype-specific differences. Of the 241 proteins found to be variable in the DA vs. 5-HT comparison, gene ontology analysis revealed enrichment of neuron- and synapse-associated cellular components and biological processes (**Fig. 6C**). In contrast to the large, stable core proteome, this pattern indicates that subtype-specific variation is concentrated in proteins associated with cytoskeletal organization and synaptic function. Together with the conserved core-proteome analysis, these results show that low-input, aspiration-based sampling can resolve both a broadly shared somal proteome backbone and distinct, sparsely overlapping differential protein subsets across neuronal subtypes.

## CONCLUSIONS

This work establishes an aspiration-based route to proteome-level phenotyping of single, identified neurons directly in intact brain tissue. By integrating patch-clamp microsampling with high-efficiency CE–ES and timsTOF MS, the approach enables targeted, *in situ* protein measurements from electrophysiologically accessed and/or fluorescently labeled somal material without prior cell isolation or dissociation. Even though only a small fraction of the somal proteome was aspirated and ∼0.25% of the processed digest was analyzed per run, the platform reproducibly quantified >1,000 proteins per neuron and resolved neuronal subtypes on the basis of protein expression alone. These results establish that controlled partial somal sampling is sufficient for proteome-driven phenotyping in intact brain tissue.

The workflow should be interpreted as aspiration-based patch proteomics rather than whole-soma retrieval. It prioritizes controlled microsampling, spatial specificity, and compatibility with optional *in situ* electrophysiology over exhaustive proteome capture. As a result, proteins enriched in distal axonal, dendritic, synaptic, or specialized membrane compartments may be underrepresented. In parallel, recent whole-soma retrieval studies have demonstrated that larger physical collections can further increase proteome depth, but they also introduce different experimental tradeoffs related to sampling completeness, spatial specificity, and coupling between recorded physiology and recovered material. These approaches are therefore complementary rather than directly equivalent and should be interpreted in the context of sampling strategy, amount of collected material, and analytical platform.

Importantly, the proteome depth achieved here was obtained on a legacy timsTOF Pro mass spectrometer. This result indicates that aspiration-based *in situ* proteotyping can reach the >1,000-protein regime without reliance on next-generation instrumentation, provided that sampling and acquisition are carefully optimized. Because the workflow is compatible with standard acute-slice preparations and a broadly accessible timsTOF Pro platform, it provides a practical foundation for extending in situ single-neuron proteomics to comparative studies of defined neuronal populations across diverse systems. Continued improvements in internal solution chemistry, sample handling, and acquisition strategies should further expand coverage and throughput, positioning in situ single-neuron proteomics as a complementary modality to transcriptomic and electrophysiological approaches for defining molecular differences among neuronal populations in native tissue.

## Acknowledgments

Parts of this work were sponsored by the Arnold and Mabel Beckman Foundation (Beckman Young Investigator award to P.N.) or the National Institute on Aging of the National Institutes of Health (R01AG088147 to P.N.).

