## Supplementary material for "Proteome-Driven Phenotyping of Identified Single Neurons in Intact Brain Tissue by Aspiration Patch Proteomics": SI Document

**Electronic Supplementary Information (SI) for**

#### SI EXPERIMENTAL

**Chemicals and Materials**. All chemicals were sourced at reagent grade or higher. Dithiothreitol (DTT), iodoacetamide (IAD), tris-hydrochloric acid (tris-HCl), tris-hydroxymethylamino-methane (Tris-base), potassium chloride (KCl), and sodium hydroxide (NaOH) were from Sigma-Aldrich (St. Louis, MO). LC-MS grade (Optima) acetic acid (AcOH), acetonitrile (ACN), formic acid (FA), and methanol (MeOH), and reagent-grade ethylenediamine tetraacetic acid (EDTA) were from Fisher Scientific (Fair Lawn, NJ). Ammonium bicarbonate (AmBIC) was from Avantor (Center Valley, PA). Sodiumdodecylsulfate (SDS) was supplied by Amresco (Solon, OH). Fused silica capillaries (40/90 µm inner/outer diameter) were from Polymicro Technologies (Phoenix, AZ) and used as received. All standards were prepared in 500-µL LoBind protein microtubes from Eppendorf (Hauppage, NY) to reduce nonspecific peptide losses on vial surfaces.

**Animals and Preparation of Brain Tissue Sections.** All protocols ensuring the humane maintenance, treatment, and use of male and female mice were approved by the Institutional Animal Care and Use Committee of the University of Maryland, College Park (approval no. R-FEB-23-09) or the George Washington University (approval no. A378). The mice were group-housed with littermates in ventilated cages in temperature- and humidity-controlled rooms with *ad libitum* access to water and rodent standard chow on a 12 h light/dark cycle.

***For method development***, wild type female and male mice were used between 9–11 weeks of age (C57BL/6, Strain 664, The Jackson Laboratory).

***For neuron phenotyping***, transgenic female and male mice were used between 9–11 weeks of age: Female and male ePet-cre (C57BL/6 background, Strain 12712, The Jackson Laboratory) and PV-cre (C57BL/6 background, Strain 8069, The Jackson Laboratory) were crossed with Ai14 (C57BL/6 background, Strain 7914, The Jackson Laboratory), hereafter ePet-cretdTomato and PV-cretdTomato, respectively, and Pitx3-eGFP (gift from Dr. Kevin Wickman).

**Single-Neuron Identification and Sampling.**

***For method development***, mice were anesthetized with isoflurane (4%) and the brain was rapidly removed and placed in ice-cold N-methyl-D-glucamine (NMDG). Horizontal brain slices (250 µm thick) containing the OB were obtained using a vibratome (Leica VT1200, Leica Biosystems Inc., IL). The brain slices were incubated at 35 ºC for 5 minutes in a holding chamber filled with NMDG. Afterwards, the slices were transferred to a second holding chamber containing artificial cerebrospinal fluid (aCSF, in mM): 125 NaCl, 2.5 KCl, 1.25 $NaH_2PO_4$, 1 $MgCl_2 \times 6 H_2O$, 11 glucose, 26 $NaHCO_3$, 2.4 $CaCl_2$, pH 7.4, and osmolarity of 290−310 mOsm at room temperature. All solutions were saturated with 95% $O_2$ and 5% $CO_2$.

OB slices were transferred to the recording chamber and perfused at a rate of 1.5 to 2.0 mL/min mounted on an Olympus BX51 W1 microscope (Evident Scientific, MA). Mitral cells were visualized using 40X objectives and DIC illumination, then patched and held at –60 mV in whole-cell patch-clamp configuration using a Sutter integrated patch clamp amplifier (1 kHz low-pass Bessel filter and 10 kHz digitization) with Igor-pro 8.04 software (Sutter Instruments).

Glass patch pipettes with resistance 2–3 MOhms were filled with internal solution containing 50 mM ammonium bicarbonate, $NH_4HCO_3$ (AmBic), which we previously optimized for patch-clamp proteomics[1]. For current clamp recordings, pipettes were filled with an internal solution of the following composition (in mM): 120 K-gluconate, 10 Na-gluconate, 4 NaCl, 10 HEPES-K, 10 Na phosphocreatine, 2 Na-ATP, 4 Mg-ATP, and 0.3 GTP, adjusted to pH 7.3 with KOH. For single-cell proteome profiling, the cytoplasm of the patched neuron was aspirated into the recording pipette immediately after entering in whole-cell configuration. Sample harvesting was completed within 3 min. Whole-cell access resistance (15-25 MΩ) and stability of the Giga-Ohm seal between the neuron and the pipette was constantly monitored to avoid contamination from the extracellular medium. The content of the pipette tip containing the harvested cytoplasm was then expelled into a 500 µL microtube.

***For neuron phenotyping***, mice were anesthetized with ketamine and dexmeditomidine (100 and 0.25 mg/kg, respectively) then perfused transcardially with NMDG-based slicing solution containing (in mM): 92 NMDG, 20 HEPES, 25 glucose, 30 $NaHCO_3$, 1.2 $NaH_2PO_4$, 2.5 KCl, 5 sodium ascorbate, 3 sodium pyruvate, 2 thiourea, 10 $MgSO_4$, and 0.5 $CaCl_2$, pH 7.4, and osmolarity of 303−308 mOsm. The brains were quickly removed and placed in ice-cold NMDG solution. Horizontal brain slices (250 µm thick) containing the dorsal raphe nucleus (DRN) or the ventral tegmental area (VTA) or coronal slices containing the prefrontal cortex (PFC) were obtained using a vibratome (Leica VT1200, Leica Biosystems Inc., IL). The brain slices were incubated at 32 ºC for 1 h in a holding chamber filled with a recovery solution containing (in mM): 92 NaCl, 20 HEPES, 25 glucose, 30 $NaHCO_3$, 1.2 $NaH_2PO_4$, 2.5 KCl, 5 sodium ascorbate, 3 sodium pyruvate, 2 thiourea, 1 $MgSO_4$, and 2 $CaCl_2$ (pH 7.4, 303−308 mOsm). Afterwards, the holding chamber containing the slices was allowed to reach room temperature. In sequence, a single slice was transferred to a chamber perfused at a rate of 1.5 to 2.0 mL/min with artificial cerebrospinal fluid (aCSF, in mM): 125 NaCl, 2.5 KCl, 1.25 $NaH_2PO_4$, 1 $MgCl_2 \times 6\, H_2O$, 11 glucose, 26 $NaHCO_3$, 2.4 $CaCl_2$, pH 7.4, and osmolarity of 303−308 mOsm at 32ºC. All solutions were saturated with 95% $O_2$ and 5% $CO_2$.

Neurons were visualized with video-assisted infrared differential interference contrast (IR-DIC) imaging. Fluorescent neurons, tdTomato-positive neurons in the DRN (ePet-cretdTomato) or in the PFC (PV-cretdTomato) and GFP-positive neurons (PitX3GFP) in the VTA, were identified by epifluorescence imaging under a water immersion objective (40×, 0.8 numerical aperture) using a Nikon Eclipse FN1 upright microscope.

tdTomato-positive and GFP-positive neurons were patched and held at –70 mV in whole-cell patch-clamp configuration using a Sutter integrated patch clamp amplifier (1 kHz low-pass Bessel filter and 10 kHz digitization) with Igor-pro 8.04 software (Sutter Instruments). Glass patch pipettes with resistance 2–3 MOhms were filled with internal solution containing 50 mM AmBic. For single-cell proteome profiling, the cytoplasm of the patched neuron was aspirated into the recording pipette immediately after entering in whole-cell configuration. Sample harvesting was completed within 3 min. Whole-cell access resistance (15-25 MΩ) and stability of the Giga-Ohm seal between the neuron and the pipette was constantly monitored to avoid contamination from the extracellular medium. The content of the pipette tip containing the harvested cytoplasm was then expelled into a 500 µL microtube.

**Bottom-Up Proteomics.** The aspirates were processed for bottom-up proteomics. Following our recent protocols to improve detection sensitivity,[1,2] the standard bottom-up workflow steps of reduction/alkylation were eliminated. Each collected protein extract was collected into 5 µL of 50 mM AmBic containing 0.1 µg of trypsin protease for one-step digestion at 60 °C for 1 h. The resulting single-neuron protein digests were vacuum-dried and stored at −80 °C until analysis. Once validated prior to our collection, each neuronal soma was collected in a 500 µL LoBind microtube with 0.1 µg trypsin protease for 1 h digestion at 60 °C. This quick digestion step enables full digestion of proteins in the sample while preventing evaporation and subsequent degradation of our already limited sample due to prolonged exposure to heat. After digestion, the samples were vacuum dried and stored at –80 °C until CE-HRMS analysis.

**CE-HRMS Proteomics.** The resulting peptides were analyzed on a custom-built, micro-loading CE platform that we recently reported[3,4]. The separation CE capillary was coaxially fed into a platinum emitter, which served as an electrospray (ESI) emitter (250/750 µm inner/outer diameter) with ~10–15 µm of CE protrusion. Peptide concentration was quantified by absorbance at 205 nm, after which an ~20 nL, containing <1 ng of protein digest, was separated by capillary zone electrophoresis in a 100-cm long capillary (40/110 µm inner/outer diameter) at ~220 V/cm field strength. The electrophoretically separated peptides were ionized in a custom-built CE electrospray ionization (ESI) ion source. In this design, the sheath solution consisting of 50% MeOH in 1% (v/v) FA was supplied through the grounded metal blunt tip emitter. The emitter was positioned ~2 mm in front of a mass spectrometer, where the spray was maintained in the cone-jet regime for efficient ion generation under guidance by a camera equipped with a long-working distance objective.[5]

For single-soma proteomes, ions with peptide-like isotope distribution were detected between *m/z* 100 to 1,700 on a trapped ion mobility spectrometer (tims) quadrupole orthogonal acceleration time-of-flight (TOF) mass spectrometer equipped with parallel accumulation serial fragmentation (PASEF) technology (timsTOF Pro, Bruker Daltonics, Billerica, MA, USA) for sequencing in a collision-induced dissociation (CID) cell. The mass spectrometer was tuned and calibrated according to vendor specifications and operated at 50,000 FWHM resolution. MS conditions were optimized for sensitivity using a pooled neuron digest. $MS^2$ was governed by data-dependent acquisition with the following settings: data acquisition rate, 2 Hz for $MS^1$ and 1 Hz for $MS^2$; survey scan cycle time, 3 s; fragmentation preference, top most-intense; $MS^2$ threshold, 250 counts per 1,000 summations; active exclusion, exclude after 1 spectra and release after 0.5 min; charge state preference, 2–5; exclude singly and unknown; *m/z* window and CID energy, 2 Da and 20–70 eV depending on charge state; collision gas, nitrogen; and dynamic exclusion, applied; smart exclusion, applied with 5× threshold; number of PASEF $MS^2$ scans, 5–25; charge range, 0–5; scheduling target intensity, 10,000–40,000; scheduling intensity threshold, 2,500 counts; collision energy and gas, 38–45 eV in nitrogen; active exclusion, reconsider precursor if current/previous intensity, 4; mobilogram, summation widths, 25 pts; max number of peaks, 3; TIMS enabled, on.

**Data Analysis.**

***Mass Spectrometry.*** The primary MS data were analyzed in FragPipe 23.1 executing MSFragger 4.3[6] against the Mus musculus proteome database (downloaded from UniProt on July 8th, 2025). The standard LFQ-MBR workflow was employed for IM-MS type data. Samples were analyzed with DDA+ enabled to allow for identification of co-isolated precursors. Peptides are reported with <1% false discovery rate (FDR) calculated against a reversed-sequence decoy database. Common contaminants were identified from the cRAP database and removed from the proteins identified in this study. Proteins that are reported in this work exclude common contaminants and proteins with a total protein score of less than 0.9.

***Statistics.*** MaxLFQ values from quantified proteins were median-normalized and $\log_{10}$-transformed. Proteins with >70% values missing across biological replicates were filtered and missing values were estimated by limit of detection (1/5 of minimum). Batch correction was performed using the ComBat method[7]. Mean maxLFQ intensity values across technical replicates were analyzed using MetaboAnalyst 6.0[8] for one-factor statistical analysis and hierarchical clustering analysis. Volcano plots were generated in MetaboAnalyst executing a Wilcoxon rank-sum test with an FDR-adjusted significance cutoff of $p = 0.05$. Fold-change thresholds were set to 1 + (3 × coefficient of variation). Hierarchical cluster analysis (HCA) was performed using Euclidean distance measurement and ward clustering based on the top 250 proteins ranked by ANOVA.

***STRING Protein–Protein Interactions.*** Stably-expressed proteins were analyzed using STRING 2025 protein-protein interaction analysis[9]. Gene IDs were searched against the *Mus musculus* genome to identify the subnetwork of proteins with documented physical interactions. Disconnected proteins were discarded and the network was analyzed by legacy k-means clustering (k = 10) to identify relevant categories of stably-expressed proteins.

**Safety Considerations.** Fused silica capillaries and borosilicate capillary emitters, which pose potential needle-stick hazard, were carefully handled. Standard safety protocols were practiced when handling chemicals. Each electrically conductive part of the CE-ESI-MS interface was grounded or isolated to prevent electrical shock.

### SI FIGURES

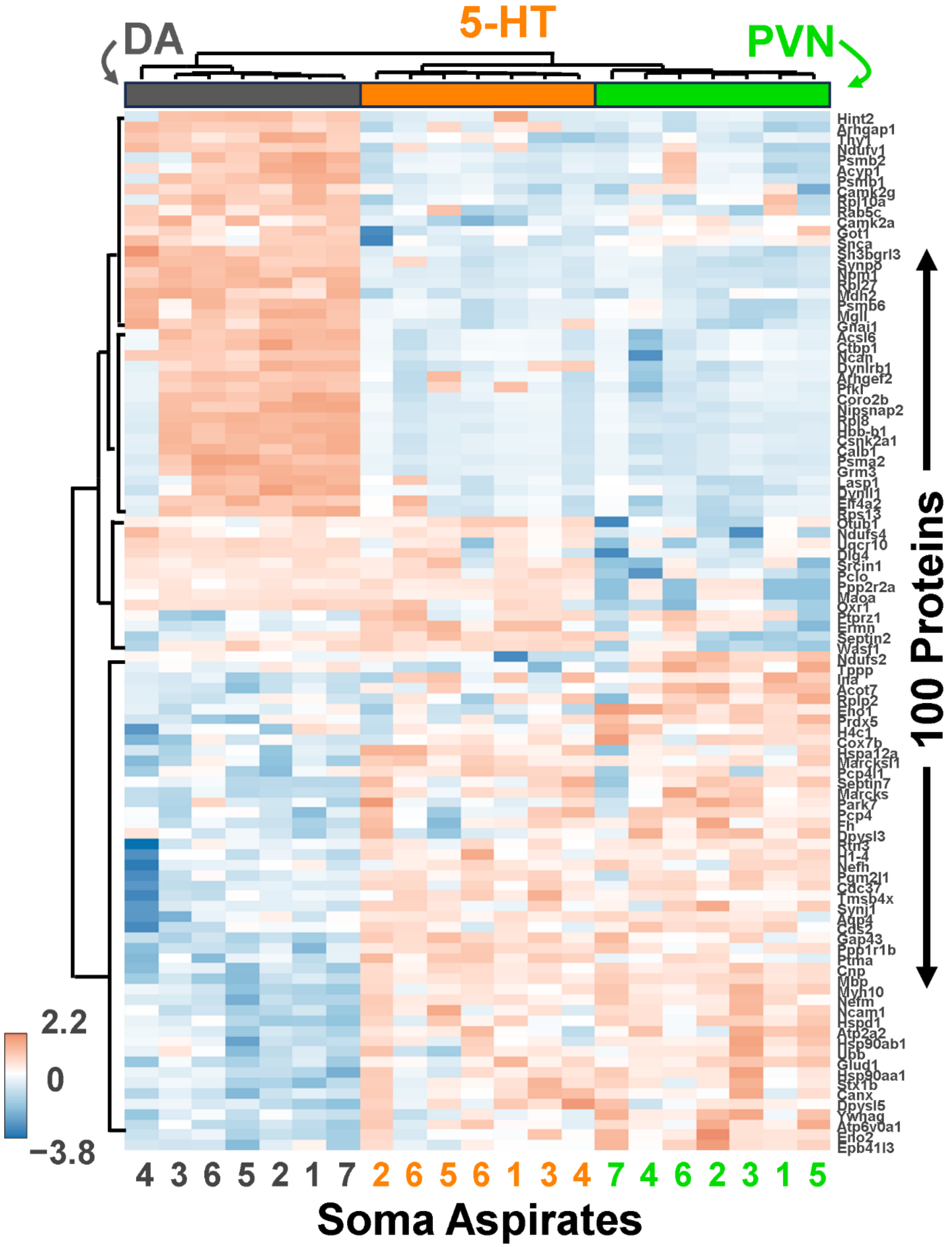

**Figure S1.** Close-up of HCA plot from **Figure 5B**. Gene names are shown.

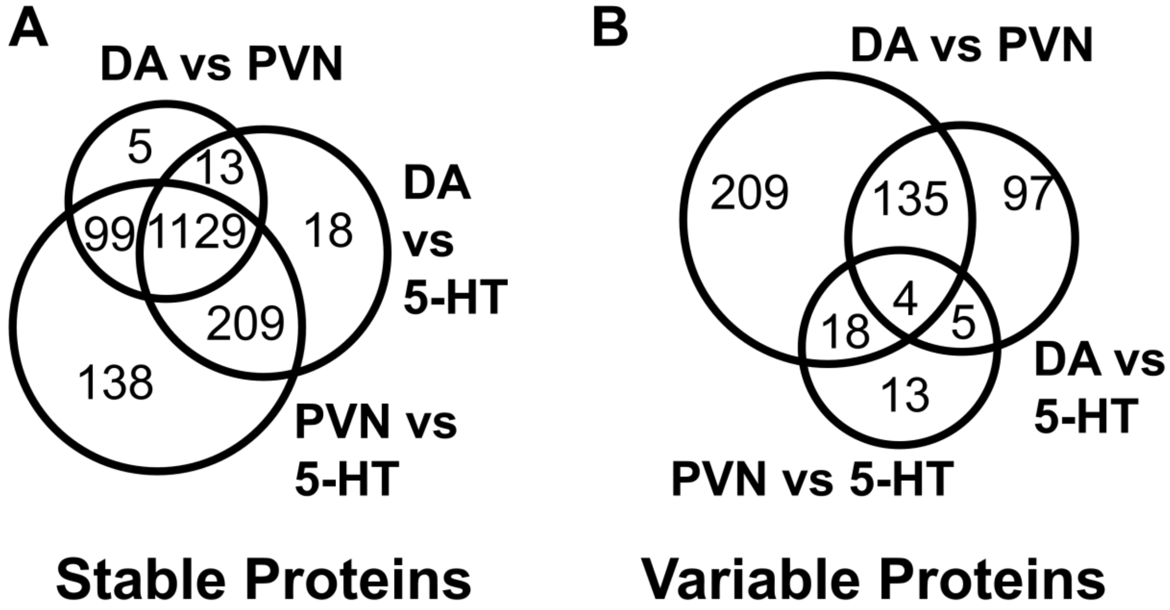

**Figure S2.** Venn diagram grouping of protein profiles among the neuronal somas with **(A)** stable and **(B)** variable expression.

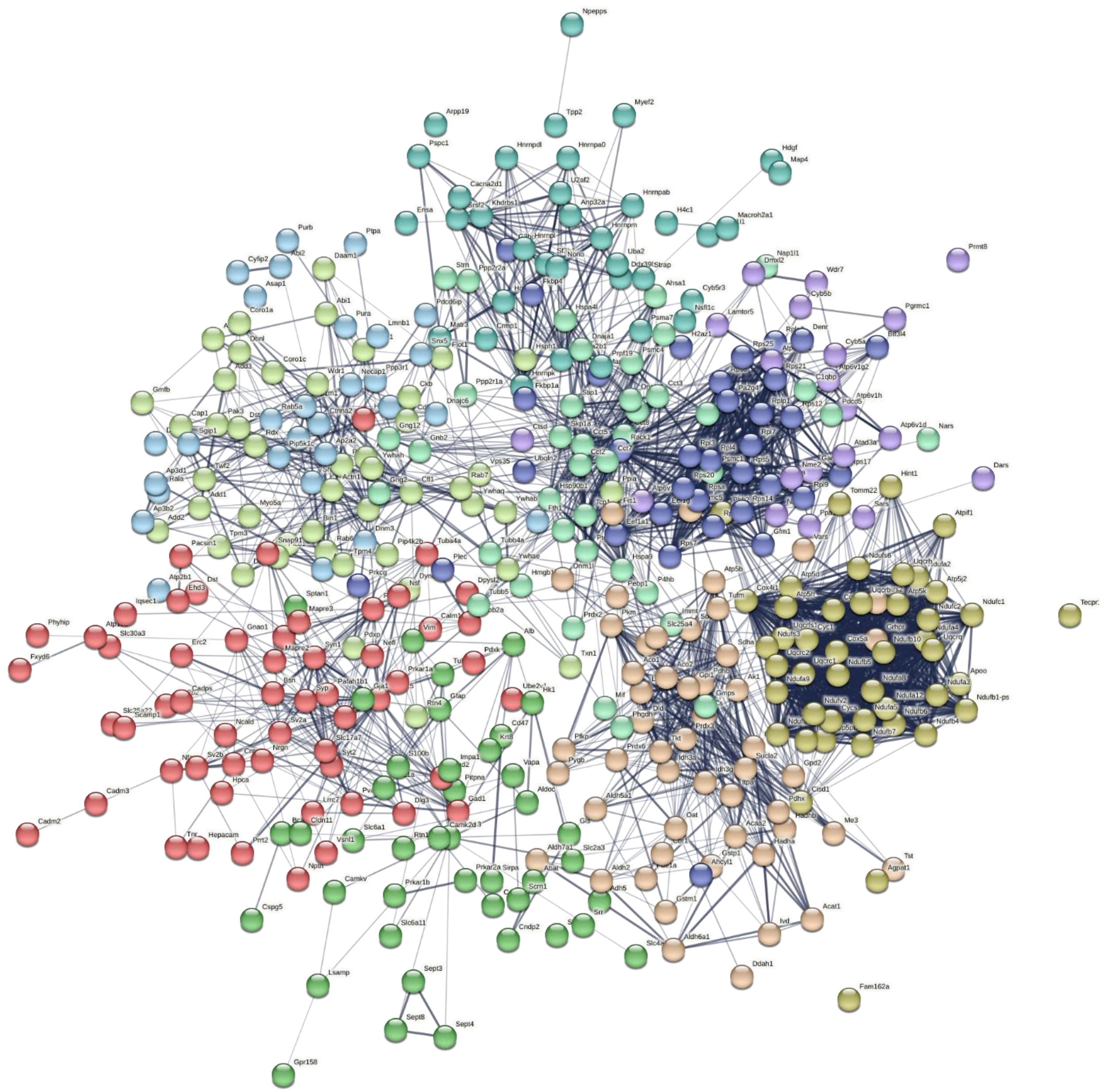

**Figure S3.** STRING analysis of the proteins that were comparable among all the neuron types ($p \geq 0.05$). The proteins are grouped into 10 clusters using K-means clustering. Network edges indicate confidence. Gene names are shown. Disconnected nodes are hidden.
